# Simultaneous PET and molecular MR imaging of cardiopulmonary fibrosis in a mouse model of left ventricular dysfunction

**DOI:** 10.1101/2023.12.15.571959

**Authors:** Brianna F. Moon, Iris Y. Zhou, Yingying Ning, Yin-Ching I. Chen, Mariane Le Fur, Sergey Shuvaev, Eman A. Akam, Hua Ma, Cesar Molinos Solsona, Jonah Weigand-Whittier, Nicholas Rotile, Lida P. Hariri, Matthew Drummond, Avery T. Boice, Samantha E. Zygmont, Yamini Sharma, Rod R. Warburton, Gregory L. Martin, Robert M. Blanton, Barry L. Fanburg, Nicholas S. Hill, Peter Caravan, Krishna C. Penumatsa

## Abstract

**Background:** Aging-associated left ventricular (LV) dysfunction promotes cardiopulmonary fibrogenic remodeling, Group 2 pulmonary hypertension (PH), and right ventricular failure. At the time of diagnosis, cardiac function has declined, and cardiopulmonary fibrosis has often developed. Here, we sought to develop a molecular positron emission tomography (PET)-magnetic resonance imaging (MRI) protocol to detect both cardiopulmonary fibrosis and fibrotic disease activity in an LV dysfunction model.

**Methods:** LV dysfunction was induced by transverse aortic constriction (TAC) in 6-month-old senescence-accelerated prone (SAMP8) mice, a subset of mice received sham surgery. Three weeks post-surgery, mice underwent simultaneous PET-MR imaging at 4.7 T. Collagen–targeted PET and fibrogenesis MR probes were intravenously administered. PET signal was computed as myocardium- or lung-to-muscle ratio (MMR, LMR). Percent signal increase (%SI) and ΔLMR were computed from the pre-/post-injection MR images. Tissue specimens were analyzed for hydroxyproline (Hyp) and allysine content. Ventricular structure and function were measured by echocardiography and hemodynamic pressure-volume (PV) loop analysis.

**Results:** Allysine in the heart (22±5 (TAC), 13±5 (sham) nmol/g, *P=0.02*) and lungs (7.5±4 (TAC), 5.8±2 (sham) nmol/lung, *P=0.17*) of TAC mice corresponded to an increase in myocardial MRI %SI (29±15 (TAC), 6.1±4 (sham), *P<0.0001*) and ΔLMR (0.3±0.1 (TAC), 0.08±0.1 (sham), *P<0.0001*). Hyp in the heart (555±90 (TAC), 400±80 (sham) µg/g, *P<0.0001*) and lungs (189±63 (TAC), 143±31 (sham) µg/lung, *P<0.01*) were elevated in TAC mice, which corresponded to an increase in PET signal (MMR: 1.8±0.1 (TAC), 1.6±0.2 (sham), *P=0.02*; LMR: 1.5±0.1 (TAC), 1.2±0.1 (sham), *P<0.001*). PV loop and echocardiography demonstrated adverse LV remodeling, function, and increased right ventricular systolic pressure in TAC mice.

**Conclusions:** Administration of collagen-targeted PET and allysine-targeted MR probes led to elevated PET-MRI signals in the myocardium and lungs of TAC mice. The study demonstrates the potential to detect fibrosis and fibrogenesis in cardiopulmonary disease through a dual molecular PET-MR imaging protocol.

## Introduction

Left ventricular (LV) compliance declines with age, which may contribute to an increased risk of heart failure in the elderly.^1,2^ Pulmonary hypertension (PH), defined as an elevation of blood pressure in the pulmonary artery,^3,4^ is a common comorbidity of left ventricular dysfunction leading to right ventricular failure,^5,6^ cardiopulmonary remodeling,^7^ and fibrosis.^8,9^

Multiple tests aid in diagnosing cardiopulmonary disease severity and progression including echocardiogram,^10,11^ magnetic resonance imaging,^12,13^ and right heart catheterization.^14^ However, at the time of diagnosis, the disease has often progressed to an advanced stage, where cardiac function has declined, and cardiopulmonary fibrosis has developed.^15,16^ It remains a challenge to detect the early onset of cardiopulmonary fibrosis and measure disease activity/fibrogenesis. The ability to diagnose LV dysfunction and PH at an early stage of the disease would allow for earlier initiation of therapy, and potentially improved prognosis.^17,18^

Tissue fibrosis and stiffness, including that of the pulmonary vasculature and the heart, occur with PH,^19,20^ cardiac overload syndromes,^21^ and aging.^19,20^ In response to elevated systemic arterial pressure, both the heart and lungs compensate to maintain homeostasis and subsequently undergo tissue remodeling, where myofibroblasts become activated and secrete inflammatory mediators and synthesize extracellular matrix (ECM) components, including collagen proteins.^22^ During this process of fibrogenesis, lysyl oxidases (LOX) are upregulated and oxidize collagen lysine residues to the aldehyde allysine which then undergoes condensation reactions with other collagen side chains to ultimately result in stable cross-links. An excess amount of ECM and ECM cross-linking leads to tissue stiffness and can ultimately lead to heart failure.^23^

We previously reported a type I collagen-targeted PET probe, ^68^Ga-CBP8, and showed that ^68^Ga-CBP8 PET could detect and stage pulmonary fibrosis in different mouse models of lung injury and in patients with idiopathic pulmonary fibrosis.^19,20,24,25^ Collagen increases with increasing disease and molecular imaging of collagen provides an overall assessment of fibrotic burden. We also showed that fibrogenesis (fibrotic disease activity), independent of overall fibrotic burden, could be noninvasively measured using a molecular MR probe targeted to allysine in animal models of lung, liver, and kidney fibrosis.^26–34^ Allysine is present at high concentrations during fibrogenesis but is rapidly consumed by cross-linking reactions once the fibrosis-causing insult is removed. Gd-1,4 is a recently described molecular MR probe that targets pairs of allysine residues and has improved sensitivity and dynamic range compared to earlier allysine-targeted probes.^29^

The advent of simultaneous PET-MRI allows 3D imaging of deep tissue structures to provide combined molecular, functional, and anatomical information. We hypothesized that cardiac and pulmonary tissue remodeling could both be quantitatively assessed through a single imaging session using appropriate molecular probes. Here, we sought to develop a dual molecular PET-MRI protocol to detect both cardiopulmonary fibrosis and fibrotic disease activity in an LV dysfunction model using collagen-targeted ^68^Ga-CBP8 PET and allysine-targeted Gd-1,4 enhanced MRI in a single imaging session. To validate cardiopulmonary tissue remodeling, we conducted tissue characterization studies including histology, immunohistochemistry (IHC), quantitative RT-PCR, and biochemical assays. LV structure and biventricular function were assessed by echocardiography and pressure-volume (PV) loop.

## Methods

### Data Availability

The data that support the findings of this study are available upon request from the corresponding authors.

### Study design

Six-to seven-month-old aging mouse models, senescence-accelerated prone mice (SAMP8) were used in this study. Left ventricular dysfunction was induced by left thoracotomy transverse aortic constriction surgery (TAC) using a 25-guage needle in SAMP8 mice (Figure 1A) as described previously.^35^ The TAC surgical procedure is provided in Supplemental Material. Control mice received sham surgery. Mice were assigned to each study group (sham, TAC) by simple randomization. All animal experiments were handled in accordance with US National Institutes of Health’s *Guide for the Care and Use of Laboratory Animals* and in compliance with the Animal Research: Reporting of In Vivo Experiments (ARRIVE) guidelines.^36^ All procedures were approved by the Tufts Medical Center and Massachusetts General Hospital (MGH) Institutional Animal Care and Use Committees. A total of eighty-one sham (n=10 female, n=33 male) and TAC (n=6 female, n=32 male) mice were used in this study. The experimental design flow chart and exclusion criteria are provided in Figure S1. All experimental evaluations were performed in a nonblinded fashion except for the histopathological and echocardiography analysis. Sample sizes and statistical results are provided in the figure legends.

**Figure 1.**
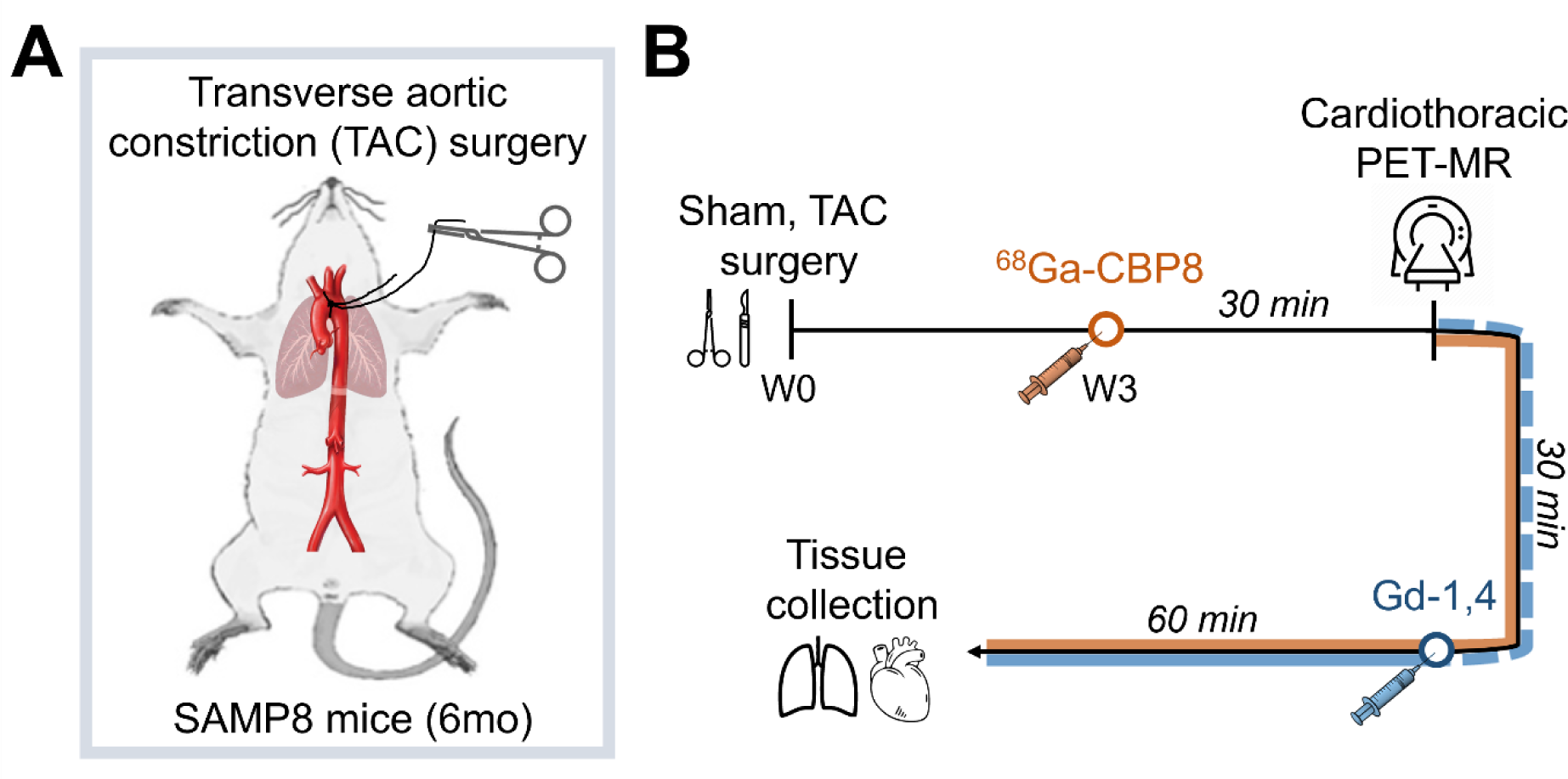
Study Design. **A**, TAC (n=6 female, n=13 male) and sham (n=7 female, n=9 male) mice underwent cardiothoracic PET-MR imaging 3 weeks post-surgery. **B**, ^68^Ga-CBP8 was intravenously administered with an injected dose between 18-34 MBq at a volume of 100-150 µL prior to the mouse being placed in the PET-MR scanner. PET (solid orange line) and pre-injection MRI (dashed blue line) scans were acquired for 30 min followed by administration of Gd-1,4 (100 µmol/kg, 75-110 µL). Cardiothoracic MRI sequences (2D T1-FLASH and 3D UTE) were repeated for a period of 60 min post-injection (solid blue line). Afterward, each mouse was euthanized, and lung and heart tissue were collected for analysis.

### Pressure-volume (PV) loop measurements

At 3 weeks post-surgery, invasive PV loop measurements were recorded in isoflurane (1% to 1.5%) anesthetized mice in supine position as previously described.^35^ Briefly, a pressure−volume (PV) catheter (Millar Instruments; Houston, Texas) was inserted into the LV (via carotid artery and aorta (pre-constriction), sham (n=3 male), TAC (n=3 male)) or RV (via jugular vein and right atrium, sham (n=6 male), TAC (n=4 male)) using a closed chest approach as described previously.^21^ PV loop data were analyzed using a LabChart application (AD Instruments, Dunedin, New Zealand). Animals were then euthanized using pentobarbital injection (120 mg/kg ip), and lungs and the hearts were inflated for histological preparation.

### Echocardiography

Cardiac function and architecture were assessed using transthoracic echocardiography on anesthetized mice (sham (n=3 male), TAC (n=4 male)) using a Vevo2100 equipped with 9-18 MHz transducer (Visualsonics Inc., Toronto, Ontario, Canada). Once the LV was clearly visualized, LV end-systolic and end-diastolic dimensions (M-mode) were measured, and the LV fractional shortening percentage and ejection fraction were calculated as previously described.^35^ LV geometry was assessed using LV weights, LV posterior wall thickness and volume during diastole. Blinded investigators performed echocardiography assessments on the day before surgery and 3 weeks post-surgery.

### Molecular Probes

^68^Ga-CBP8 is a peptide-based PET probe targeting type I collagen, with a dissociation constant (K_d_) of 2.1 ± 0.1 μM for type I human collagen and 4.6 ± 0.5 μM for rat collagen.^19^ ^68^Ga-CBP8 was synthesized as previously reported^19,20,25,37^ with some modifications described in Supplemental Material. Gd-1,4 is a dual binding fibrogenesis probe targeting allysine, with a relaxivity of 17.4 ± 1.0 mM^-1^s^-1^ when bound to allysine containing protein (BSA^Ald^) at 1.41 T, 37 °C, and was synthesized as described.^29^

### Cardiothoracic PET-MRI

Mice (sham (n=7 female, n=9 male), TAC (n=6 female, n=13 male)) were imaged 3 weeks post-surgery on a 4.7 Tesla Bruker Biospec MRI equipped with a PET insert (Bruker, Billerica, MA). Mice were anesthetized with isoflurane (1 to 2%). The body temperature was maintained at 37°C, and inhaled isoflurane concentration was adjusted to maintain a respiration rate of 60±5 breaths per minute. The tail vein was cannulated for intravenous delivery of molecular probes.

^68^Ga-CBP8 was intravenously administered with an injected dose between 18-34 MBq at a volume of 100-150 µL prior to the mouse being placed in the PET-MR scanner. Next, the mouse was placed in the scanner, and PET-MRI was acquired simultaneously (Figure 1B). Pre-injection MRI scans were acquired for 30 min and encompassed cardiac/respiratory gated 2D T1-weighted fast low-angle shot (FLASH): repetition time (TR)/echo time (TE)/ inversion time (TI) = 609.5/3.5/590 ms, number of averages (NEX) = 3, bandwidth (BW) = 390.6 Hz/pixel, flip angle (FA) = 50°, field of view (FOV) = 40 × 40 mm^2^, matrix size = 200 × 200, slice thickness = 1 mm, acquisition time (AT) = 6:05 min; 3D ultrashort echo time (UTE): TR/TE = 4/0.012 ms, NEX = 1, BW = 976.6 Hz/pixel, FA = 16°, FOV = 32 × 32 × 32 mm^3^, matrix size = 128 × 128 × 128, AT = 3:25 min; and 3D FLASH: TR/TE = 15/2.5 ms, NEX = 4, BW = 1250 Hz/pixel, FA = 12°, FOV = 27.5 × 48 × 24 mm^3^, matrix size = 60 × 120 × 60, AT = 3:36 min. Then Gd-1,4 was administered (100 µmol/kg, 75-110 µL) followed by 2D FLASH and 3D UTE MRI which were repeated for a period of 60 min.

After the PET-MR imaging session each mouse was euthanized followed by tissue collection. The left lung (LL), right lung (RL), muscle, liver, kidney, blood, and tail were collected from all animals. Additionally, the heart was separated into three parts: right ventricle (RV), septum (sep), and LV. All tissue samples were weighed, followed by measuring the radioactivity with a gamma counter (Wizard2Auto Gamma, PerkinElmer). Biodistribution measurements were calculated based on injected dose (ID), decay-corrected, and presented as %ID/g for all organs. Lung measurements were reported as %ID/lung.

### PET Image Analysis

The PET datasets were reconstructed using maximum *a* posteriori probability (MAP 0.5 mm) algorithm run over 10 iterations to a voxel size of 0.5 × 0.5 × 0.5 mm^3^. PET data were reconstructed in 6 × 5 min frames and 3 × 10 min frames. Radioactivity measurements were performed on averaged PET frames from 60-90 min post-injection.

Reconstructed PET-MR data were quantitatively evaluated using AMIDE software package.^38^ MR images (3D FLASH and UTE) were used to draw volumes of interest (VOIs) over the following organs: LL, RL, myocardium (septal and anterior wall of the left ventricle). An additional two VOIs were drawn on the left and right forelimb muscles. The VOIs were applied to decay-corrected PET scans. Radioactivity measurements were converted to counts per milliliter per minute and then divided by the injected dose (ID) to obtain an imaging VOI-derived percentage of the injected radioactive dose per cubic centimeter (%ID/cc). PET measurements were also reported as the ratio of tissue signal (lung or myocardium) to muscle (LMR, MMR).

### Magnetic Resonance Image Analysis

For cardiac 2D FLASH images, a region of interest (ROI) was manually drawn in ImageJ (Fiji, version 1.0) and encompassed the septal and anterior wall of the left ventricle while avoiding the blood pool. A second ROI was placed on the phantom doped with Gd-DOTA to normalize the signal intensity across all the 2D FLASH images. Myocardial signal intensity (SI) was measured in each normalized 2D FLASH image. SI from pre-injection images (n=3) were averaged. %SI increase was calculated by subtracting the pre-injection SI from the post-injection SI then dividing by the pre-injection SI (Eq. 1).

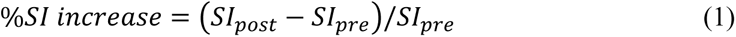

Pulmonary 3D UTE images were normalized to the signal intensity of a Gd-DOTA doped phantom placed next to the animals during image acquisition. VOI measurements included LL, RL, and muscle. LMR was calculated by dividing the SI in the lungs by the SI of the forelimb triceps muscle (Eq. 2) as performed in previous pulmonary disease studies.^27,34^ LMR from pre-injection scans (n=3) were averaged. LMR from RL and LL VOIs were averaged for each UTE image. ΔLMR was calculated by subtracting the pre-injection LMR from the post-injection LMR (Eq. 3). In addition, pulmonary %SI increase was computed from 3D UTE images using (Eq. 1).

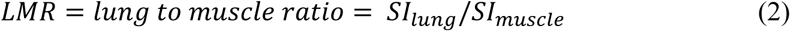

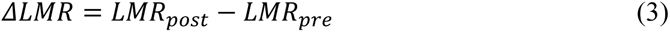

%SI increase and ΔLMR were averaged 20-45 min post-injection of the Gd-1,4 probe, and dynamic curves are shown in Supplemental Material. Change in pulmonary tissue to muscle contrast to noise ratio (ΔCNR) was computed from the pre- and post-injection MR images as described in Supplemental Material.

### Histology and immunohistochemistry

A subset of cardiac (sham (n=5 male), TAC (n=8 male)) and pulmonary (sham (n=4 male), TAC (n=6 male)) tissue samples was fixed with 4% formaldehyde, paraffin-embedded, and sliced at 5 µm thickness. Cardiac samples were cut in the short axis at the mid ventricle and lung samples were inflated prior to embedding. Trichrome staining was performed by the Specialized Histopathology Services – MGH Core using standard methods.

LOX and TG2 protein expressions in the lung and heart tissues were detected using immunohistochemistry (IHC) assays with antibodies against LOX (1:250; NB100-2527, Novus Biologicals) and TG2 (1:250; ab421, Abcam); protocol provided as Supplemental Material.

The slides were scanned using NanoZoomer Slide Scanner (Hamamatsu) at x20 original magnification. For cardiac tissue samples, the percentage of fibrosis was measured from Trichome slides using ImageJ (Fiji, version 1.0) with a uniform color threshold segmentation approach.^39,40^ The percent fibrosis was computed with respect to the area of the LV. For pulmonary tissue samples, the regions of fibrosis and disease classification (mild, moderate, severe) were determined by an experienced pulmonary pathologist (L.P.H.).

### RNA isolation and quantitative RT-PCR

Quantitative RT-PCR was performed to measure the gene expression in collagen, type 1, alpha 1 (*Col1a1*) and lysyl oxidase (*LOX-1*, *LOX-2*, *LOX-3*, *LOX-4*) as demonstrated in previous mouse models^33^. To isolate RNA and conduct quantitative RT-PCR, frozen heart and lung tissues were utilized (sham (n=8 male), TAC (n=8 male)). The RNA extraction was performed using TRIzol (Invitrogen) according to the manufacturer’s instructions. In summary, 1µg of extracted RNA was reverse transcribed using a High-Capacity cDNA Reverse Transcription Kit.

During the real-time PCR, the Applied Biosystems TaqMan Gene Expression Master Mix was used with predeveloped primer/probe assays from Thermo Fisher Scientific for *Col1a1* (Mm0080166-g1), *LOX-1* (Mm 00495386_m1), *LOX-2* (Mm00804739_m1), *LOX-3* (Mm 01184865_m1), *LOX-4* (Mm 00446385_m1) and 18S ribosomal RNA primer/probe set (Hs 03003631; Thermo Fisher Scientific).

For Fibronectin, Real-time polymerase chain reaction (PCR) analysis was performed using 2x SYBR Green Master Mix (Thermo Fisher Scientific) on an ABI Prism 7900 Sequence Detection System (Thermo Fisher Scientific) as described previously.^41^ Human and mouse specific primer sets (IDT Technologies, Coralville, IA) were used.

To normalize the cycle threshold (Ct) values, 18s ribosomal RNA levels were used. The relative quantification of specific genes was determined using the ΔΔCt method as previously described.^41,42^

### Biochemical determination of hydroxyproline and allysine

Hydroxyproline (Hyp) in the LV (sham (n=10 female, n=14 male), TAC (n=6 female, n=12 male)) and RL (sham (n=10 female, n=14 male), TAC (n=6 female, n=12 male)) was measured with high-performance liquid chromatography (HPLC) following a published method.^43^ Allysine of the septum (sham (n=3 female, n=2 male), TAC (n=3 female, n=2 male)) and LL (sham (n=9 female, n=13 male), TAC (n=6 female, n=12 male)) was quantified using HPLC following a published method.^34,44^ Measurements are reported as amounts per wet weight of tissue [g] or amounts per lung.

### Statistical Analysis

Quantitative measurements are displayed as box plots. The bottom and top of the box represent the first and third quartiles, the center band is the mean, whiskers report the minimum and maximum values, and individual values are displayed as points on the plot. Measurements are reported as mean±SD, significant differences are reported when P<0.05. The Kolmogorov–Smirnov test was used to assess normality in each dataset and an F-test was used to compare variances. If the datasets had a normal distribution and equal variances an unpaired two-tailed t-test was used to compare differences between two groups (sham, TAC). If the datasets had a normal distribution and unequal variances an unpaired two-tailed t-test with Welch’s correction was performed. If the datasets did not have a normal distribution a Mann Whitney test was performed. All statistical analyses were performed in GraphPad Prism 9.0 (GraphPad software).

## Results

In the heart, trichrome staining showed myocardial interstitial fibrosis in TAC mice (Figure 2A), which corresponded to a significant increase in percent fibrotic area with respect to the left ventricle area compared to sham mice (1.6±0.3% (sham, n=5) vs 4.8±2.1% (TAC, n=8), *P=0.006*, Figure 2B). Hyp, a measure of total collagen and marker of fibrosis, was significantly elevated in the myocardium of TAC mice (400±80 (sham, n=24), 555±90 (TAC, n=18) µg/g, *P<0.0001*, Figure 2C). Allysine, a marker of fibrogenesis, was significantly elevated in the myocardium of TAC mice (13±6 (sham, n=5) vs 22±5 (TAC, n=5) nmol/g, *P=0.02*, Figure 2D). Gene expression of proteins associated with fibrogenesis: collagen type 1, alpha 1 (*Col1a1*) and lysyl oxidase paralogs *LOX-1, LOX-2, LOX-3,* and *LOX-4*, showed that these were all significantly elevated in the LV myocardium of TAC mice compared to sham (*P<0.05*, Figure 2E). In addition, *Col1a1* and *LOX-4* expression in the RV myocardium was significantly elevated in TAC mice (Figure S2A).

**Figure 2.**
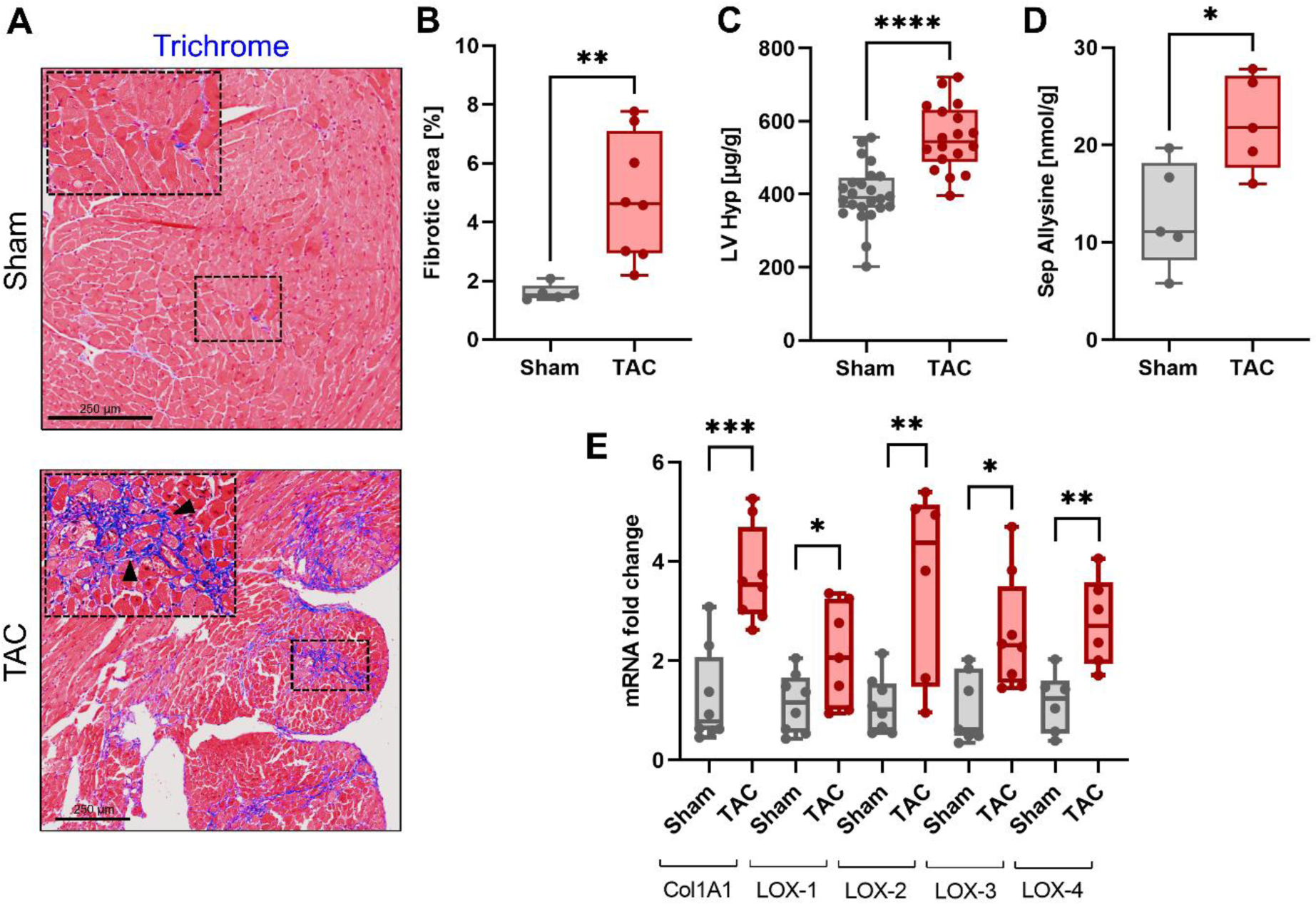
Heart tissue characterization. **A**, Representative Trichrome staining of the myocardium shows regions of interstitial fibrosis in TAC SAMP8 mice (inset, arrowheads). **B**, The percentage of IHC Trichrome fibrosis-positive tissue with respect to the LV area of the short-axis slice (n≥5 per group). **C**, Hydroxyproline content as a marker of myocardial fibrosis in sham and TAC mice (n≥13 per group). **D**, Allysine content as a marker of myocardial fibrogenesis measured in the septum (Sep) of sham and TAC mice (n=5 per group). **E**, Gene expression levels of collagen, type 1, alpha 1 (*Col1a1*) and lysyl oxidase (*LOX-1*, *LOX-2*, *LOX-3*, *LOX-4*) in sham and TAC SAMP8 mice (n≥6 per group). Fold changes were normalized with housekeeping gene (18s). All data are shown as means ± SD; *P < 0.05, **P < 0.01, and ***P < 0.001; ns, not significant; one-way ANOVA, post hoc comparison, two-tailed.

Histopathology of the pulmonary tissue showed heterogeneity in the disease response, 3/6 TAC mice had moderate to severe lung injury, but the remaining TAC mice had mild lung injury (Figures 3A). Lung injury was not detected in sham mice (Figures 3A). TAC mice with moderate to severe disease showed several histopathological findings of acute lung injury and tissue remodeling including 1) inflammation, 2) focal regions of fibrosis surrounding blood vessels and airways, 3) presence of hemosiderin and hemosiderosis, and 4) leaked fibronectin into the extracellular space. LOX and TG2 staining were present predominately around pulmonary blood vessels indicating active tissue remodeling and distribution of LOX and TG2 proteins (Figure 3A). Focal regions of fibrotic activity led to a moderate but significant elevation in Hyp of the whole RL (143±31 (sham, n=24), 189±63 (TAC, n=18) µg/lung, *P<0.01*, Figure 3B) of TAC mice compared to sham. Allysine in the whole LL (LL: 5.8±2 (sham, n=17), 7.5±4 (TAC, n=16) nmol/lung, *P=0.17*, Figure 3C) was elevated in TAC mice, but this elevation was not statistically significant. However, RT-qPCR uncovered significantly elevated *Col1a1, LOX-2, LOX-3,* and *LOX-4* gene expression in the lungs (*P<0.05*, Figure 3D). Additionally, TAC mice had significantly lower body weight (BW) with a significant elevation of RV weight to BW and lung weight to BW, compared to sham mice indicating cardiopulmonary disease progression (Figure S2B-E).

**Figure 3.**
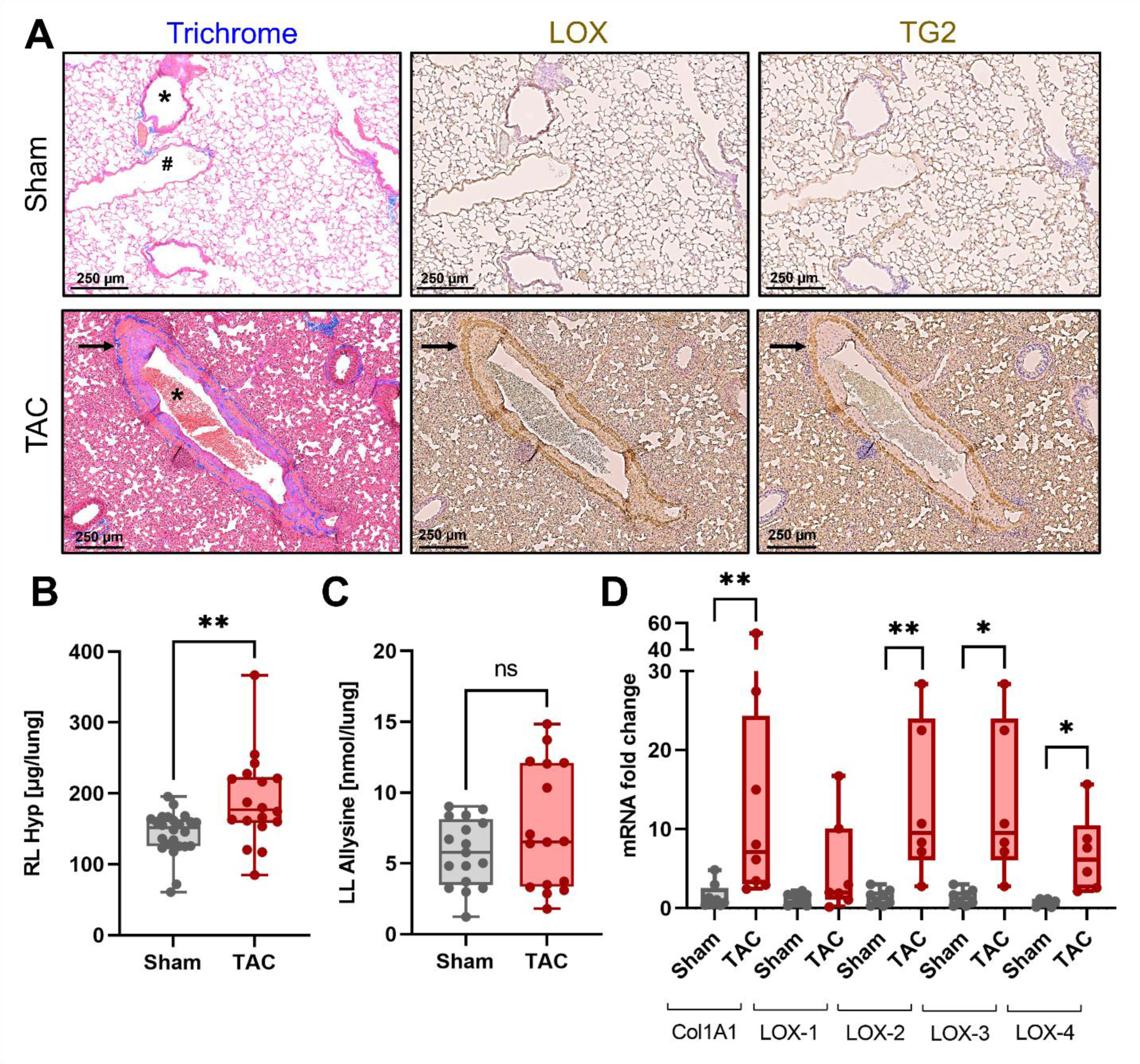
Lung tissue characterization. **A**, Representative histology staining shows vessel remodeling (arrows) through fibrosis (Trichrome), LOX-positive (LOX), and TG2-positive (TG2) regions in the lungs of TAC SAMP8 mice. Sham mice did not exhibit vessel remodeling. Pulmonary vessels (*), airways (#). **B**, Hydroxyproline content as a marker of pulmonary fibrosis measured in the right lung (RL) of sham and TAC mice (n≥12 per group). **C**, Allysine content as a marker of pulmonary fibrogenesis measured in the left lung (LL) of sham and TAC mice (n≥13 per group). **D**, Gene expression levels of collagen, type 1, alpha 1 (*Col1a1*), and lysyl oxidase (*LOX-1*, *LOX-2*, *LOX-3*, *LOX-4*) in the lungs of sham and TAC SAMP8 mice (n≥6 per group). Fold changes were normalized with housekeeping gene (18s). All data are shown as means ± SD; *P < 0.05, and **P < 0.01; ns, not significant; one-way ANOVA, post hoc comparison, two-tailed.

Echocardiographic (Table 1, Figure S3) and invasive hemodynamic PV loop (Table 2) measurements were collected to assess LV remodeling and PH secondary to LV dysfunction in the TAC mice. Echocardiographic M-mode measurements showed signs of left ventricular hypertrophy, including increases in LV weight (P=0.008), internal diameter at end-diastole (P=0.008) and end-systole (P=0.017), posterior wall thickness (P=0.037) and volume (P=0.009) at end diastole in TAC mice. In addition, echocardiographic data showed LV ejection fraction (P=0.008) and fractional shortening (P=0.006) were significantly reduced, demonstrating TAC surgery induced LV systolic dysfunction. PV loop functional analysis also identified markers of LV dysfunction including significantly elevated aortic systolic (P=0.005) and LV end diastolic (P=0.011) pressures as well as reduced aortic diastolic pressure (P=0.007), cardiac output (P=0.003), stroke volume (P=0.004) and stroke work (P=0.036) in TAC mice. Effective arterial elastance, a measure of arterial load, was significantly elevated in TAC mice (P=0.012). Right ventricular systolic pressure was significantly elevated to 42.75±12.8 mmHg in TAC mice compared to 22.8±3.65 mmHg in sham mice (P=0.006), indicative of PH in the TAC mice.

**Table 1.** Echocardiographic assessment of left ventricular structure and function. Left ventricle (LV); left ventricular internal diameter at end diastole (LVIDd) and at end systole (LVIDs); left ventricular posterior wall thickness at end diastole (LVPWd); left ventricular volume during diastole (LV Vol;d); ejection fraction (EF); fractional shortening (FS). All data are shown as means ± SD; *P < 0.05, and **P < 0.01; t-test, two-tailed.

| Measurement | Sham (n=3) | TAC (n=4) | P* |
| --- | --- | --- | --- |
| LV Weights (mg) | 125 ± 18 | 260 ± 53 | 0.009** |
| LVIDd (mm) | 3.3 ± 0.4 | 4.3 ± 0.3 | 0.008** |
| LVIDs (mm) | 2.0 ± 0.8 | 3.5 ± 0.4 | 0.017* |
| LVPWd (mm) | 1.1 ± 0.2 | 1.5 ± 0.2 | 0.037* |
| LV Vol;d (uL) | 43.7 ± 10.7 | 82.7 ± 13.7 | 0.009** |
| EF (%) | 60 ± 3 | 40 ± 8 | 0.008** |
| FS (%) | 31 ± 2 | 19 ± 4 | 0.006** |

**Table 2.**
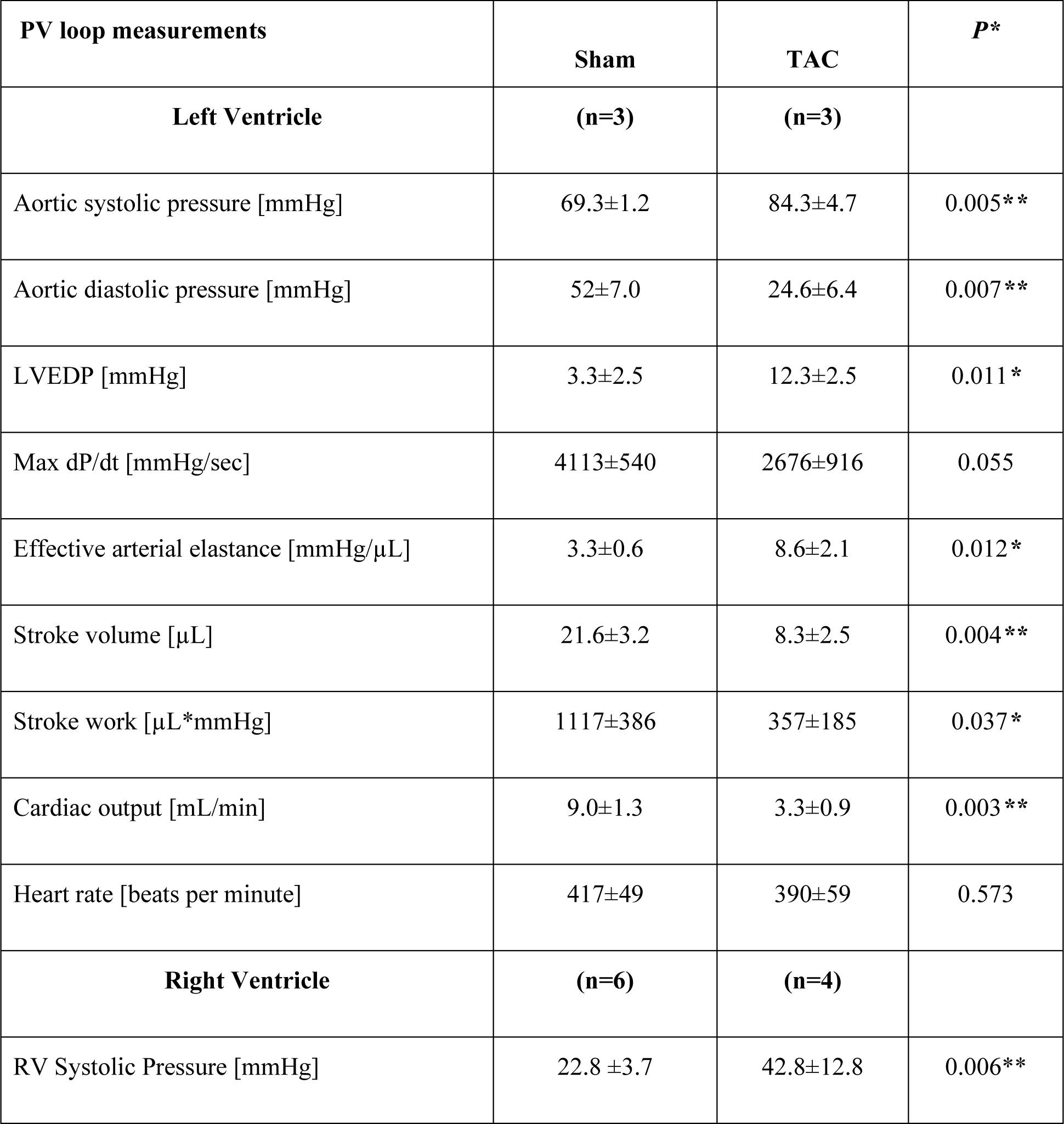
Hemodynamic pressure-volume loop measurements. PV, pressure volume; LV, left ventricle; LVEDP, left ventricular end diastolic pressure; Maximum rate of pressure change in the left ventricle, Max dP/dt; right ventricle, RV. All data are shown as means ± SD; *P < 0.05, and **P < 0.01; t-test, two-tailed.

MRI and PET were simultaneously acquired in the heart and lungs (Figure 4A,B; 5A,B; S4-7,9-10). MRI %SI increase in the myocardium was significantly elevated in TAC mice (29±15% (TAC, n=19) vs 6.1±4% (sham, n=14), *P<0.0001*, Figure 4C, S4B). ^68^Ga-CBP8 PET showed a significant increase in radioactivity in the myocardium of TAC mice (1.6±0.8 (TAC, n=10) vs 0.7±0.2 (sham, n=10) %ID/cc, *P=0.003*). In addition, the MMR PET values were significantly elevated (1.8±0.1 (TAC, n=10) vs 1.6±0.2 (sham, n=10), *P=0.02*) 65-95 min post-injection of ^68^Ga-CBP8 (Figure 4D,E, S7, S8B).

**Figure 4.**
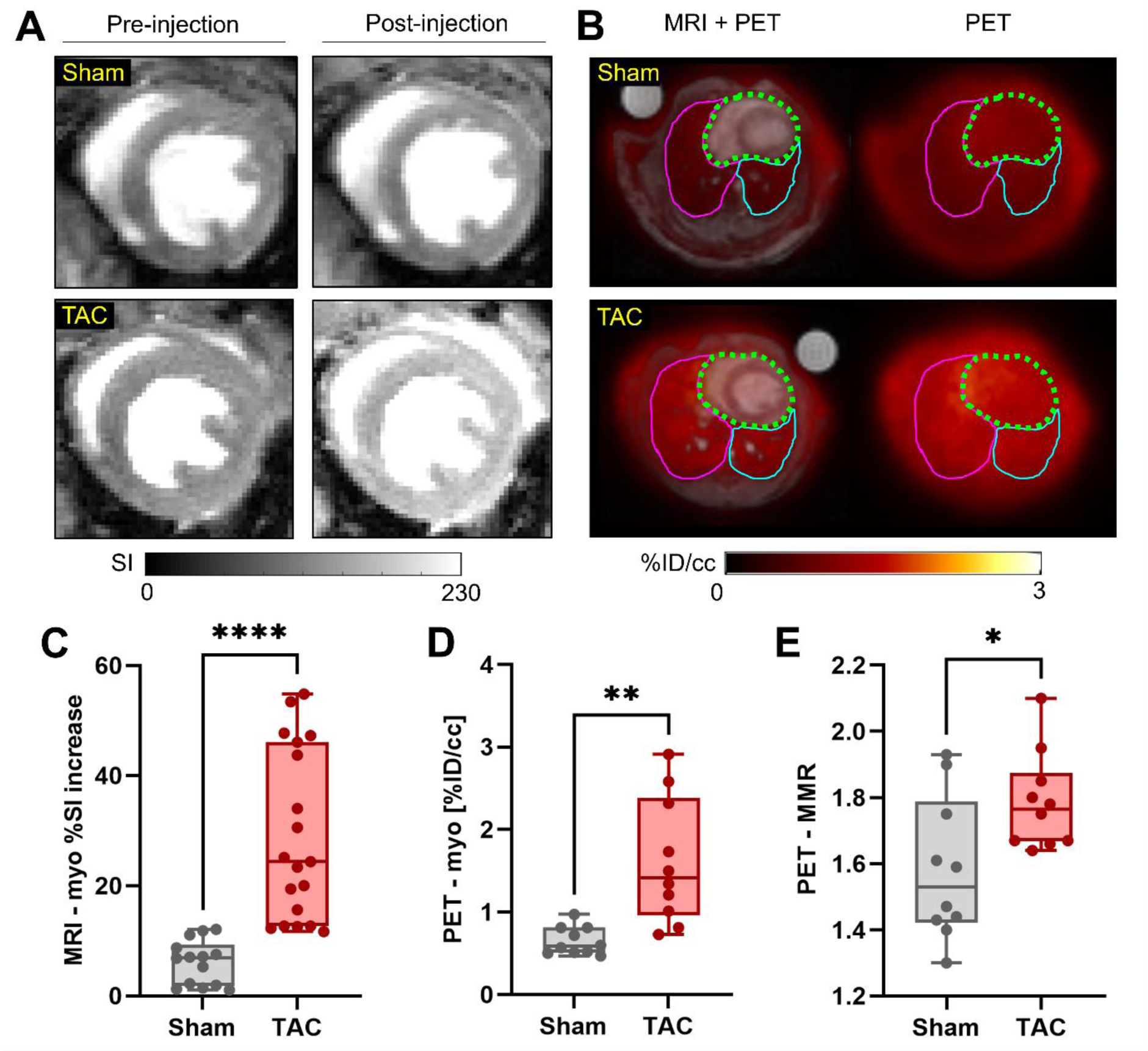
Heart Imaging. **A**, Short axis myocardial 2D T1 FLASH images from sham and TAC mice pre-injection and 20 min post-injection of Gd-1,4 (100 µmol/kg iv). **B**, Axial MRI-PET fused images and PET-only images (heart: dashed green line, RL: fuchsia solid line, LL: blue solid line) from sham and TAC mice 86 min post-injection of ^68^Ga-CBP8 (28 MBq iv). **C**, %SI increase in the myocardium relative to pre-injection images at 20-45 min post-injection of Gd-1,4 (n≥9 per group). **D**, Myocardium (myo) and **E**, myo-to-muscle PET signal 65-95 min post-injection of ^68^Ga-CBP8 (n=10 per group). All data are shown as means ± SD; *P < 0.05, **P < 0.01, and ****P < 0.0001; one-way ANOVA, post hoc comparison, two-tailed.

**Figure 5.**
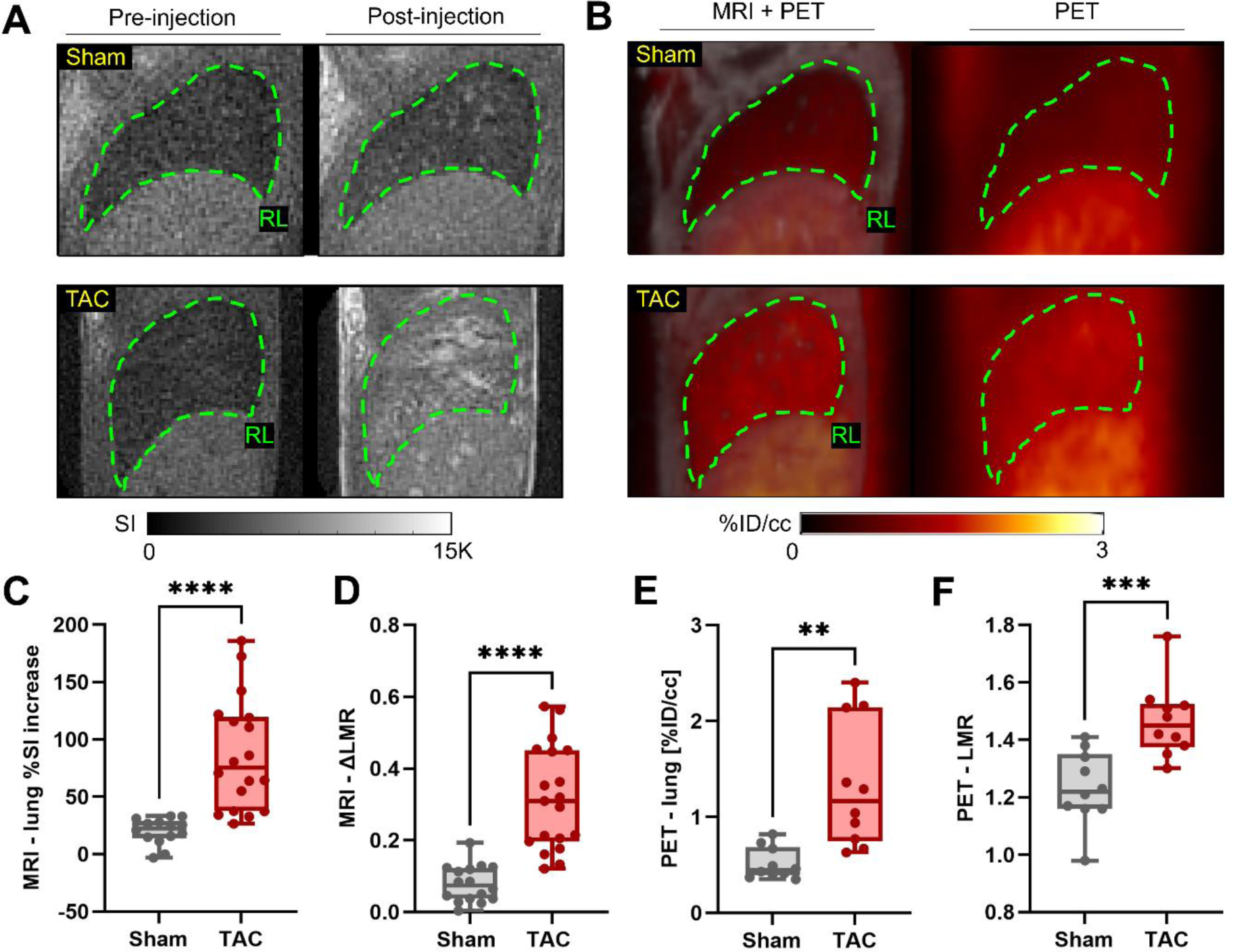
Lung Imaging. **A**, Sagittal UTE images from sham and TAC mice pre-injection and 20-45 min post-injection of Gd-1,4 (100 µmol/kg iv). **B**, Sagittal MRI+PET and PET-only images (RL: dashed green line) from sham and TAC mice 86 min post-injection of ^68^Ga-CBP8 (28 MBq iv). **C**, %SI increase and **D**, change in lung to muscle contrast-to-noise ratio (ΔCNR) relative to pre-injection images at 20-45 min post-injection of Gd-1,4 (n≥7 per group). **E**, Lung and **F**, lung-to-muscle PET signal 65-95 min post-injection of ^68^Ga-CBP8 (n=10 per group). All data are shown as means ± SD; *P < 0.05, and **P < 0.01; one-way ANOVA, post hoc comparison, two-tailed.

From UTE MR imaging of the lungs, TAC mice showed a significantly higher mean pulmonary %SI increase (87±49 (TAC, n=18) vs 20±11 (sham, n=14), *P<0.0001*) and ΔLMR (0.3±0.1 (TAC, n=18) vs 0.08±0.1 (sham, n=14), *P<0.0001*) at 20-45 min post-injection of Gd-1,4 (Figure 5C,D, S5B). In addition, the lungs of TAC mice had a significant increase in pre-injection contrast-to-noise (CNR) and LMR (Figure S6A,B). The ΔCNR from pre- and post-injection imaging was significantly elevated in the lungs of TAC mice (Figure S5C, S6C,D). ^68^Ga-CBP8 PET showed a significant increase in lung radioactivity of TAC mice (1.3±0.7 (TAC, n=10) vs 0.5±0.2 (sham, n=10) %ID/cc, *P=0.003*) and the ratio of LMR PET values were significantly increased (1.5±0.1 (TAC, n=10) vs 1.2±0.1 (sham, n=10), *P<0.001*) 65-95 min post-injection of ^68^Ga-CBP8 (Figure 5E,F, S7, S8C). There were no differences in MRI or PET measurements between RL and LL (Figure S9). Ex vivo biodistribution studies showed significantly increased activity in the heart and lungs of TAC mice (Figure S10).

## Discussion

Clinical prevalence of LV dysfunction with heart failure increases with aging.^45,46^ If left untreated, the associated elevated pressure overload in the pulmonary circulation can progress to secondary PH and RV failure with poor clinical outcomes.^8,9^ Physiologic responses to aging-related cardiac pressure overload have been linked to structure-function abnormalities associated with fibrogenic remodeling leading to decreased tissue compliance and LV dysfunction.^47,48^ However, there is a lack of sensitive noninvasive tools available to assess the extent of fibrosis and early onset of disease. Previous studies from our lab and others have shown that SAMP8, a mouse model of aging, and TAC-induced LV pressure overload mouse model developed cardiac and pulmonary fibrosis and PH associated with LV dysfunction.^8,9,41,49^

In consideration of the common pathophysiological features of aging and pressure-overload, we have now analyzed if TAC-induced pressure overload in an aging mouse model exacerbates the fibrogenic remodeling in lung and heart tissues. Here we further proposed a dual imaging modality (PET-MR) to image at a systems level (both heart and lungs) of disease in a model with pulmonary and cardiac injury using two imaging probes that targeted type 1 collagen (^68^Ga-CBP8) or allysine (Gd-1,4).

Fibrosis is a common response to injury and characterized by excess deposition of collagens, primarily type I collagen. Both the amount of fibrosis and whether the disease is ongoing are important clinical questions. Previous studies showcase that the PET probe, ^68^Ga-CBP8, and its MRI analog, EP-3533, can detect type 1 collagen and stage disease in preclinical models of pulmonary,^19,20,50^ cardiac,^51,52^ and hepatic^53–55^ fibrosis. Both probes, ^68^Ga-CBP8 and EP-3533, have been extensively validated through blocking experiments, comparison to control probes, and gain/loss of function models. The MRI probe, Gd-1-4, has its own unique target, allysine, and allysine-targeted MRI has been employed to detect fibrogenesis in models of lung, liver, and kidney fibrosis.^26–33^ Both probes, ^68^Ga-CBP8 and Gd-1,4, do not interfere with one another and were able to detect focal regions of fibrosis and disease activity in the heart and lungs.

Our studies with a more severe form of TAC (26- or 27-gauge) produced significant mortality in the 6-month-old SAMP8 mice (data not shown). More importantly, despite the milder form of TAC (25-guage) in the senescence-accelerated mice, allysine biochemical assay and LOX IHC demonstrated evidence of active fibrosis in the heart and lungs, which corresponded to imaging findings. The SAMP8 TAC mice had focal regions of pulmonary fibrosis surrounding arteries and airways, which is a characteristic of PH tissue remodeling.^56–58^ Of note, we previously reported that a pro-fibrogenic crosslinking enzyme, tissue transglutaminase (TG2), was elevated in a pressure-overloaded TAC^41^ and a senescent-prone mouse model of aging^49^ where myocardial fibrosis and HFpEF have been described. We also identified that elevation of TG2 activity is causative for fibrogenic remodeling including collagen deposition in experimental mouse models of cardiopulmonary fibrosis.^41^ Consistent with these findings, using immunohistochemical imaging, we now for the first time show that TAC surgery induced TG2 expression in both pulmonary arteries and airways (Figure 3A). Taken together, assisted by novel biochemical and imaging methods, the study further supports that SAMP8 TAC mice show signs of tissue fibrogenic remodeling in heart and lungs at an advanced stage of heart failure.

In the clinic, cardiovascular MRI techniques like late gadolinium enhancement (LGE)^59–61^ and extracellular volume (ECV) mapping^62,63^ provide an indirect measurement of collagen deposition in the heart, but do not provide an assessment of disease activity or pulmonary fibrosis. Pulmonary hypertension is diagnosed by right heart catheterization to measure pulmonary arterial pressures. Here we were able to detect both fibrosis and fibrogenesis in the heart and lungs noninvasively in a mouse model that resulted in a mild and heterogeneous disease of LV dysfunction and secondary PH. The dual molecular PET-MR imaging protocol described here has the potential to stage cardiopulmonary disease, provide early diagnosis of disease activity and to monitor treatment response.

There are some limitations to the study. The long aging period of the mice (6-month) in combination with a complex invasive surgical TAC procedure limited the number of experiments we could perform. Previous work investigated the sex differences of the TAC model and demonstrated that male mice develop increased disease severity.^64,65^ Our initial studies showed disease was more severe in male mice, and therefore, after initial experiments using both sexes, we focused the current study on male mice, which limits the generalizability of these results to females. Future studies would need to include detailed comparisons between the sexes. Secondly, we prioritized the specific-binding probes during imaging studies. Ideally, negative control probes^19^ would have been used in a subset of mice to validate that signal elevation was caused by specific-binding of the imaging probe. However, extensive work was performed to establish the specific-binding of the PET probe, ^68^Ga-CBP8, in the lungs^19,20^ and the specific-binding of the MR-probe, Gd-1,4, in the liver.^29^ Additionally, the fibrosis and fibrogenesis were validated through histology, RT-qPCR, and HPLC biochemical measurements of Hyp and allysine. Thirdly, compared to clinical cardiothoracic MR, the sequences available on the small animal scanner require much longer acquisition times which limited our imaging protocol to focus on molecular imaging. Extending this work to a human protocol would likely include established protocols to assess changes in wall thickness and cardiac function.

In conclusion, injection of collagen-targeted ^68^Ga-CBP8 and allysine-targeted Gd-1,4 probes led to elevated PET and MRI signals, respectively, which correlated with fibrosis and fibrogenesis markers. To our knowledge, this is the first study to implement a dual molecular MRI and PET imaging protocol using two molecular probes to image two organs. Overall, the study demonstrates the potential to assess disease activity and burden at a systems level by characterizing heart and lung disease during a single imaging session. Future work will include a longitudinal study to assess disease progression and therapeutic response in a mouse model of aging-associated LV dysfunction and PH.

## Abbreviations

(fibrogenesis): Active fibrosis
(ΔLMR): Change in lung to muscle MRI signal ratio
(CNR): Contrast-to-noise
(ΔCNR): Change in CNR
(EF): Ejection fraction
(ECM): Extracellular matrix
(FS): Fractional Shortening
(Hyp): Hydroxyproline
(LL): Left lung
(LV): Left ventricle
(LVEDP): LV end-diastolic pressure
(LVIDd): Left ventricular internal diameter at end diastole
(LVIDs): Left ventricular internal diameter at end systole
(LVPWd): Left ventricular posterior wall thickness
(LOX): Lysyl oxidases
(UTE): MRI sequence: 3D ultrashort echo time
(FLASH): MRI sequence: T1-weighted fast low-angle shot
(%ID/cc): Percent injected dose per cubic centimeter of tissue
(%SI): Percent signal increase
(PH): Pulmonary hypertension
(LMR): Ratio of lung to muscle
(MMR): Ratio of myocardium to muscle
(RL): Right lung
(RV): Right ventricle
(RVSP): RV systolic pressure
(PV) Loop: Pressure-volume
(SAMP8): Senescence-accelerated prone
(TG2): Transglutaminase
(TAC): Transverse aortic constriction
(CBP8): Type I collagen–targeted PET probe
(UTE): Ultrashort echo time

## Affiliations

Department of Radiology, Athinoula A. Martinos Center for Biomedical Imaging, Massachusetts General Hospital and Harvard Medical School, Boston, MA, United States (**B.F.M., I.Y.Z., Y.N., Y.I.C., M.L.F., S.S., H.M., J.W.W., N.R., A.T.B., S.E.Z., P.C.**). Institute for Innovation in Imaging, Massachusetts General Hospital, Boston, MA, United States (**B.F.M., I.Y.Z., Y.N., M.L.F., S.S., H.M., N.R., A.T.B., S.E.Z., P.C.**). Department of Medicine, Division of Cardiology, Massachusetts General Hospital and Harvard Medical School, Boston, MA, United States (**E.A.A.**). Bruker BioSpin, Preclinical Imaging, Billerica, MA, United States (**C.M.S.**). Department of Medicine, Division of Pulmonary and Critical Care Medicine, Massachusetts General Hospital and Harvard Medical School, Boston, MA, United States (**L.P.H., M.D.**). Department of Pathology, Massachusetts General Hospital, Harvard Medical School, Boston, MA, United States (**L.P.H.**). Division of Pulmonary, Critical Care and Sleep Medicine, Tufts Medical Center, Boston, MA, United States (**Y.S., R.R.W., B.L.F., N.S.H., K.C.P.**). Molecular Cardiology Research Institute, Tufts Medical Center, Boston, MA, United States (**G.L.M.**, **R.M.B.**).

## Acknowledgements

We greatly appreciate support from the Specialized Histopathology Services - MGH Core for histology staining.

## Author Contributions

Study concept and design, **B.F.M., I.Y.Z., B.L.F., N.S.H., K.C.P., P.C.**; left thoracotomy transverse aortic constriction surgery, **N.S.H., G.L.M., R.M.B., R.R.W.**; pressure-volume loop experiments, **G.L.M., R.M.B., R.R.W., K.C.P.**; echocardiography experiments, **R.M.B., R.R.W., K.C.P.**; development and synthesis of molecular fibrogenesis MR-probe (Gd-1,4); **Y.N., E.A.A.**; radiolabeling of type I collagen binding PET-probe (^68^Ga-CBP8), **M.L.F., S.S., H.M. S.E.Z.**; animal preparation for imaging studies and probe injections **B.F.M., J.W.W., N.R.**; cardiothoracic PET-MRI acquisition and image processing, **B.F.M., I.Y.Z. Y.I.C., C.M.S. J.W.W., N.R.**; tissue collection, **B.F.M., R.R.W., M.D., A.T.B.**; biodistribution measurements, **B.F.M., M.D., A.T.B.**; histology and immunohistochemistry, **L.P.H., B.F.M., R.R.W., Y.S., K.C.P.**; RNA isolation and quantitative RT-PCR, **R.R.W., Y.S., K.C.P.**; biochemical assays, **B.F.M., M.L.F., S.S., E.A.A., H.M., M.D.**; statistical analysis, **all authors**; manuscript drafting and editing for important intellectual content, **all authors**; and approval of final version of submitted manuscript, **all authors**.

## Supplemental Material

Supplemental Methods Figures S1 – S10

